# The R2T(P) complex orchestrates the SHQ1 driven early steps of box H/ACA snoRNP maturation

**DOI:** 10.64898/2026.05.30.726169

**Authors:** Philipp Busse, Ana C. F. Paiva, Paulo E. Santo, Hugo Gizardin-Fredon, Carolina P. Cassona, Marie-Eve Chagot, Catarina F. Malta, Céline Verheggen, Hannah Oksannen, Sarah j. Butcher, Sarah Cianferani, Xavier Manival, Celia Plisson-Chastang, Edouard Bertrand, Pedro M. F. Sousa, Tiago M. Bandeiras

**Author notes:** Department of Chemistry-Biochemistry, Biocenter II, Johannes Gutenberg University Mainz, 55128 Mainz, Germany.

## Abstract

The chaperones RuvBL1 and RuvBL2 are members of the AAA+ ATPase family and participate in diverse cellular processes, including DNA repair, transcriptional regulation, and assembly of macromolecular complexes such as snoRNPs. The biogenesis of box H/ACA snoRNPs additionally requires the assembly factor SHQ1. These protein-RNA complexes are essential for ribosome biogenesis and telomerase stability and are linked to diseases such as dyskeratosis congenita and cancer. Despite detailed knowledge of mature complexes, their assembly mechanisms and how they can be modulated remain unclear.

We characterize a trimeric interaction between SHQ1 and RuvBL1:RuvBL2, providing insight into early maturation of the protein-only precursor of box H/ACA snoRNPs. SHQ1 binds the flexible domain II of RuvBL1:RuvBL2, corresponding to the dodecamerization interface, suggesting disruption of this interface and promotion of hexamer formation.

We further purified a complex containing RuvBL1:RuvBL2, SHQ1, and DKC1, the catalytic component of H/ACA snoRNPs and a client of SHQ1. This demonstrates that SHQ1 and DKC1 can simultaneously associate with RuvBL1:RuvBL2, potentially facilitating DKC1 release and subsequent snoRNA binding. Additionally, we identified a direct interaction between SHQ1 and RPAP3, a co-chaperone of RuvBL1:RuvBL2 (within the R2TP). Hence, RPAP3 may be responsible for recruiting SHQ1:DKC1 to hexameric RuvBL1:RuvBL2 and/or assist in the AAA+ mediated dissociation of the dimer. Since SHQ1 shares a domain with PIH1D1, an integral member of the R2TP complex, our findings suggest that early H/ACA snoRNP maturation may involve the R2T instead of the previously proposed R2TP complex.

## Introduction

The biological function of all molecules is determined by their native three-dimensional shape, which usually marks the lowest energetic state a functional protein can adapt to [1]. Large molecules or those that contain hydrophobic patches typically require assistance during folding to circumvent the clustering of kinetic traps, which can ultimately result in protein misfolding or aggregation [2, 3]. Kinetic traps and protein aggregates mark a non-functional state of low folding energy [1]. Chaperones assist during their client’s folding process to shift the equilibrium between protein folding and unfolding toward natively folded and functional states, thereby ensuring cellular homeostasis [3–5]. Additionally, chaperones that are members of the AAA+ ATPase family can utilize the energy of ATP hydrolysis to undergo conformational changes, which generate mechanical forces that either actively fold proteins [5] or sequester protein aggregates, enabling iterative cycles of client folding [6]. Some chaperones are essential for cell survival, underscoring the importance of chaperone-assisted protein folding [7]. Malfunctional chaperones have been linked to cancer development [8, 9]; hence, an in-depth understanding of how they impact various cellular pathways is crucial for the development of novel therapies.

Besides facilitating protein domain folding, some chaperones guide the assembly of multi-protein complexes. RuvBL1 (R1) and RuvBL2 (R2) are highly conserved AAA+ ATPases that form homo- and heterohexameric rings as well as heterododecameric double rings and assist numerous DNA-related functions like transcriptional regulation or chromatin maintenance [10, 11]. Among AAA+ ATPases, they are unique as they bear a flexible arm-like domain called domain II (DII), which is inserted inside domain I’s primary structure. All DIIs protrude at the same side of the R1R2 ring, dividing the complex into two structural units: the rigid AAA+ core module formed by DI and DIII, and the flexible DIIs [12]. The flexible arms act as regulatory elements, mediating client interactions, and serve as the dimerization interface between two R1R2 rings to form dodecamers, a possible R1R2 storage form.

ATP hydrolysis triggers a conformational change that also involves the DIIs, exerting mechanical forces that might be essential for client complex assembly. The proteins PIH1D1 and RPAP3 have been identified as R1R2 interactors, forming a complex called R2TP [13, 14]. This complex, in turn, acts as a HSP90 co-chaperone, and together they play a major role in the assembly of macromolecular complexes like RNA polymerases, the Telomerase complex, and small nucleolar ribonucleoproteins (snoRNPs) [15]. Most snoRNPs modify RNAs post-transcriptionally, including the two most frequent modifications: 2’O-methylation [16] and pseudouridine isomerization [17, 18]. Based on the snoRNAs’ motifs, snoRNPs are divided into two major classes: box C/D and box H/ACA, which associate with different sets of core proteins. The snoRNA serves as a guide to identify the position to be modified in the complex’s target RNA, while the protein components stabilize the complex and catalyze the reaction. Box H/ACA snoRNPs are responsible for the isomerization of uridine to pseudo-uridine, a common modification in ribosomal RNAs (rRNAs) before they are incorporated into the ribosome [19]. This modification enables the formation of an additional hydrogen bond, increasing RNA stability.

Human box H/ACA snoRNAs are between 60 and 150 nucleotides long and can either be individually transcribed from DNA by RNA polymerase-II, or are coded within genes and are generated by mRNA splicing. The genomic DNA codes for over 150 different box H/ACA snoRNAs, which convey target variability to box H/ACA snoRNPs [20].

Being the catalytic active protein of the H/ACA’s RNA-protein complex, Dyskerin Pseudouridine Synthase 1 (DKC1) is the central component of the human RNA-protein complex. Besides its pseudouridine synthase activity [21], DKC1 plays a role in stabilizing and maintaining the telomerase complex [22]. DKC1 comprises a <u>P</u>seudo<u>U</u>ridine synthase and <u>A</u>rchaeosine transglycosylase (PUA) domain, which binds RNAs, and a catalytic tRNA pseudouridine synthase B (TruB) domain, which drives the pseudouridinylation. Mutations in the DKC1 encoding gene lead to Dyskeratosis Congenita (DC) [23], an inherited human disease, and have also been identified in several forms of cancer [24]. DKC1 is prone to aggregation [25] and may always require tight chaperoning. Hence, its isolation is challenging, and most of its structural information has resulted from the three-dimensional structure determination of DKC1-containing macromolecular complexes or from homology models of its archaeal and yeast homologues [21].

SHQ1 is a box H/ACA assembly factor, and as such, it is conserved throughout all eukaryotes. It was proposed that it binds DKC1 shortly after its translation in the cytoplasm, ensuring immediate chaperoning [26] [27]. SHQ1 harbors an N-terminal HSP90 co-chaperone p23-like CHORD and Sgt1 domain (CS) and the C-terminal Shq1-specific domain (SSD). Whereas the CS domain might bind DKC1’s amino-terminal region, the SSD domain is proposed to mimic RNA and bind to DKC1’s PUA domain [28]. The two SHQ1 domains may form a stabilizing clamp around DKC1, which prevents non-specific RNA interactions [26]. During the maturation of the protein-only pre-RNP complex, SHQ1 must dissociate from DKC1 to enable its binding to box H/ACA snoRNA [26]. As DKC1 contains a nuclear localization signal (NLS), it seems possible that the SHQ1:DKC1 complex is imported to the nucleus, where SHQ1 might dissociate and allow snoRNP maturation [29].

Dysfunctional SHQ1 is associated with neurodevelopmental and movement disorders, as mutations cause the recessively inherited diseases “dystonia-35 childhood onset” and “neurodevelopmental disorder with seizures and dystonia” (NEDDS) [30].

There is evidence that the HSP90/R2TP co-chaperone complex is involved in box H/ACA snoRNP maturation [13, 31–37], as R1R2 was identified to interact with DKC1 and SHQ1 *in vitro* [35, 38], and SHQ1 was identified to interact with RPAP3 in SILAC IP studies [39]. Here, we report the direct interaction of human SHQ1 with R1R2 complexes, as well as the biophysical and biochemical characterization of R1R2:SHQ1 complexes, including the determination of their three-dimensional structure in single-particle cryo-electron microscopy (cryo-EM). We were also able to show that RPAP3 specifically interacts with SHQ1 *in vitro*. The successful purification and biophysical characterization of R1R2:SHQ1:DKC1 complexes, together with the aforementioned R1R2:SHQ1 characterization, allowed us to propose a novel mechanism for the maturation of the protein-only precursor box H/ACA snoRNP complexes. As R1R2, SHQ1, and DKC1 are involved in various pathways crucial for cellular homeostasis, assessing their molecular interactions may also aid in designing novel drugs to treat neurodevelopmental diseases and/or cancer.

## Results

### The complex of RuvB-Like1 and RuvB-Like2 specifically interacts with SHQ1

The human protein SHQ1 homologue (SHQ1) was proposed to mimic RNA in its interaction with Dyskerin (DKC1) [28], and the complex consisting of the AAA+ ATPases RuvB-Like1 (R1) and RuvB-Like2 (R2) is known to be crucial for box H/ACA snoRNP biogenesis [40]. We therefore evaluated whether SHQ1 and the R1R2 complex were directly interacting.

To rule out mediated interaction through a third partner, as it would be possible in the previously published pull-down experiments [26], we expressed and purified recombinant and histidine (His)-tagged SHQ1 from *E. coli* cells. Purified SHQ1 was loaded onto a preparative Superdex S200 26/60 size exclusion chromatography (SEC) column and eluted as an apparent dimer (~130 kDa) (data not shown). The identity of SHQ1 was confirmed by western blotting against its His-tag and by peptide mapping by mass spectrometry (Supplementary Figure 1), which also allowed us to determine the purity of the purified protein to be 99 %. Using nanoDSF, we determined the melting temperature of SHQ1 to be 49.2 °C (Supplementary Figure 2), indicating that it is a sufficiently stable protein. However, we could further stabilize the protein by +1.5 °C by changing the buffer formulation from Tris to phosphate.

The stability of SHQ1 allowed us to assess whether it interacts with R1R2 surfaces (whose purification we previously established [41]) in SPR experiments (Figure 1A). While we detected a specific interaction between SHQ1 and R1R2 complexes, a 1:1 binding model interaction could not be applied. The interaction profile appears highly heterogeneous, most likely due to multiple SHQ1 binding sites within the R1R2 complex, displaying distinct interaction kinetics. Despite this heterogeneity, the interaction affinity was calculated to be 0.94 nM, suggesting that R1R2 and SHQ1 interact with a very high affinity.

**Figure 1:**
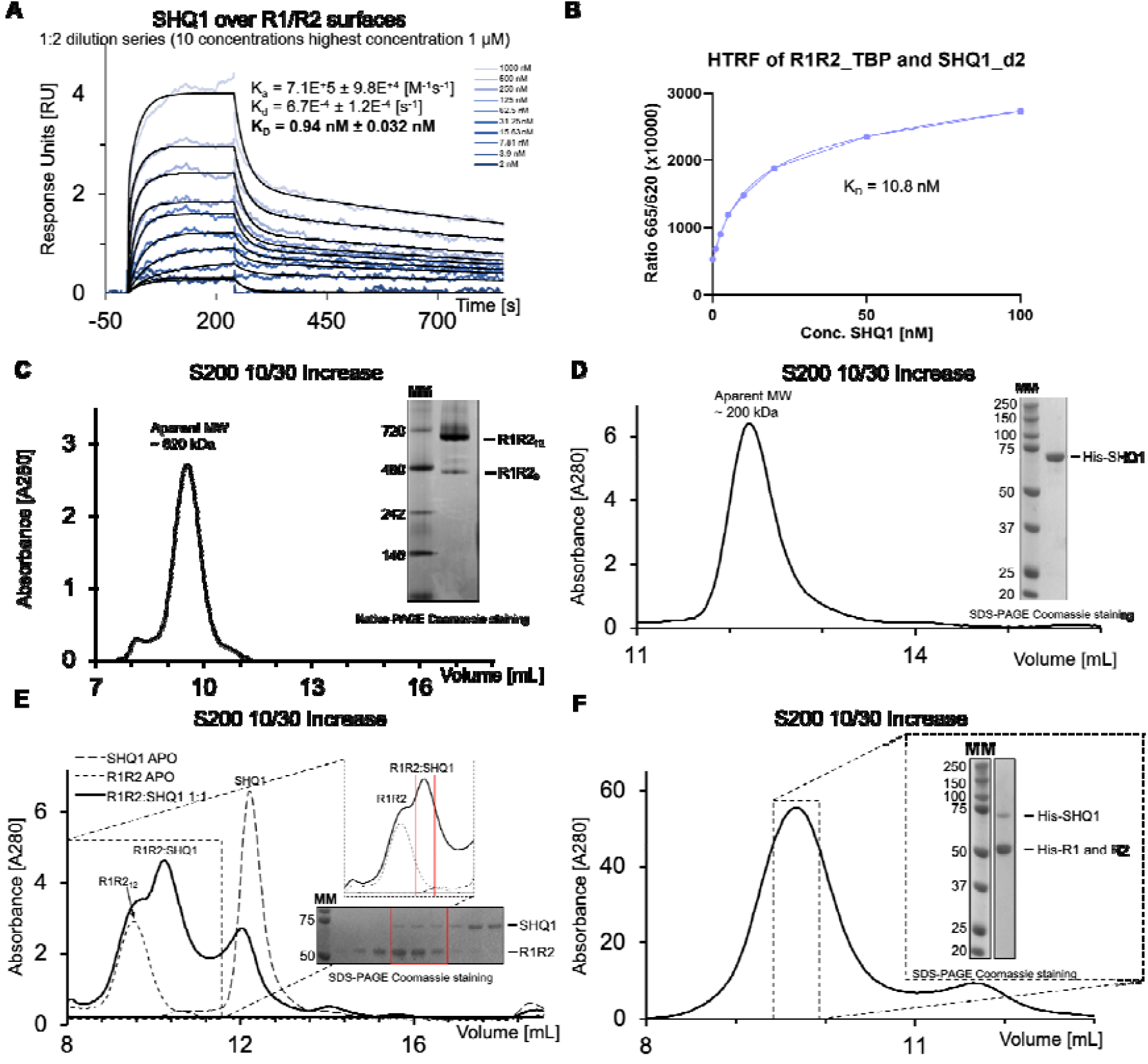
The RuvBL1:RuvBL2 complex interacts with SHQ1. **A:** SPR experiment: The R1/R2 complex has been immobilized on CM5 sensor chips via standard amine coupling. SHQ1 was injected at 10 different concentrations generated by a 1:2 dilution with 5 µM representing the highest concentration. The concentration-dependent increase in the SPR signal shows the direct interaction between the R1R2 complex and SHQ1. The dissociation constant was calculated to be 0.94 nM. **B:** Confirmation of the detected interaction by HTRF. To record the fluorescence transfer, 1 nM of R1R2_TBP, assuming their hexameric form, and 1-100 nM of SHQ1_d2 were used. All data points were recorded in triplicates, multiplied with 10.000, and are presented with their respective standard deviations. A binding constant similar to that obtained in the SPR experiment was calculated: 10.8 nM. **C:** Chromatogram of isolated R1R2 complex loaded onto an S200 10/300 size exclusion chromatography column. The elution volume corresponds to a complex of roughly 620 kDa. However, the complexes elute close to the columns’ void volume, exceeding its optimal separation size. The inserted native PAGE shows that most of the complex migrates below the 720 kDa marker, suggesting dodecameric (620 kDa) R1R2 complexes. Additionally, it shows a faint band below the 480 kDa marker, representing the hexameric (310 kDa) fraction of isolated R1R2. **D:** Chromatogram of purified His-SHQ1 loaded onto an S200 10/300 size exclusion chromatography column. The elution volume corresponds to roughly 200 kDa, indicating SHQ1 (65 kDa) may form trimers (195 kDa). The inserted SDS-PAGE shows the purified His-SHQ1 with no visible contamination of other proteins. **E:** Superimposition of chromatograms showing isolated R1R2 complex (dotted line), isolated His-SHQ1 (dashed line), and their 1:1 molar mixture (continuous black line) loaded onto an S200 10/300 size exclusion chromatography column. Zooming into the chromatogram proves the appearance of a new peak shifted towards a later elution compared with the isolated R1R2 complex. The inserted SDS-PAGE shows the chromatographic fractions, and the red box highlights those fractions corresponding to the additional peak, embodying the co-elution of R1R2 and SHQ1. **F:** Re-injection of the R1R2:SHQ1 peak fraction after mixing the proteins in a 1:6 molar ratio. The inserted SDS-PAGE proves the complex formation and its purity.

To confirm our finding via an orthogonal method, we labeled the R1R2 complexes with Terbium-Trisbipyridine (TBP), a fluorescence donor, and SHQ1 with d2, an acceptor, to perform homogeneous time-resolved fluorescence (HTRF) experiments (Figure 1B and Supplementary Figure 3). The detected fluorescence transfer and the nanomolar binding affinity (21 nM), calculated from steady-state measurements, confirmed our initial observations by SPR.

Encouraged by the identification of a high-affinity interaction, we aimed to reconstitute R1R2:SHQ1 complexes from separately purified proteins. In previous purifications, we reported the predominant R1R2 complex species to be dodecameric (~620 kDa) [41] (Figure 1C). The molecular weight of SHQ1 is approximately 65 kDa, as predicted by its amino acid composition and SDS-PAGE migration profile (Figure 1D). In solution, the oligomerization state appears to be concentration-dependent, with dimeric or trimeric species observed, as suggested by comparison of the preparative and analytical SEC profiles (Figure 1D and Supplementary Figure 4). To ensure optimal R1R2:SHQ1 complex formation, we mixed the proteins in a 1:6 (R1R2 hexamer to SHQ1 monomer) molar ratio and collected the peak corresponding to the formed complex. Incubating the proteins for 16 hours was followed by injection onto an analytical SEC column to purify any complex that had formed from the isolated proteins. The comparison between the individual proteins’ SEC profiles and those of the incubated proteins revealed the formation of an additional peak, smaller than the R1R2 dodecamers but larger than SHQ1 (Figure 1E). Analysis of the newly formed peak by SDS-PAGE with subsequent Coomassie staining revealed the co-elution of R1R2 and SHQ1 (Figure 1E insertion).

Sample homogeneity was reached by several rounds of re-injection of the collected peak fractions onto the SEC column (Figure 1F).

As the R1R2:SHQ1 complex seemed to be smaller than the R1R2 dodecamers, we hypothesized that the interaction of SHQ1 with R1R2 dodecamers may lead to their disruption into hexamers, as previously observed for other R1R2-client complexes [41].

### Determination of the R1R2:SHQ1 complex stoichiometry

In order to identify the oligomeric state of R1R2 within the purified R1R2:SHQ1, we performed mass photometry. We analyzed the R1R2 complex at 10, 25, and 50 nM (Figure 2A), which showed mass peaks fitting R1R2 dodecamers (676 ± 21 kDa), hexamers (338 ± 19kDa), and monomers (59 ± 17 kDa). The discrepancies observed between SEC (Figure 1C, only dodecamer detected) and mass photometry (Figure 2A, equilibrium between dodecamers and hexamers) might be explained by the lower protein concentration (nanomolar range) required to perform mass photometry. Equal molarities of the R1R2:SHQ1 complex, analyzed by mass photometry, revealed a fairly homogeneous sample, with the main species corresponding to ~400 kDa, which is consistent with R1R2 hexamers interacting with SHQ1 moieties and in agreement with our SEC observations. By zooming into this 400 kDa peak, several sub-populations can be detected corresponding to the binding of one (384 ± 25 kDa, 61 % abundance) and two (447 ± 31 kDa, 30 % abundance) SHQ1 molecules (Figure 2B and C).

**Figure 2:**
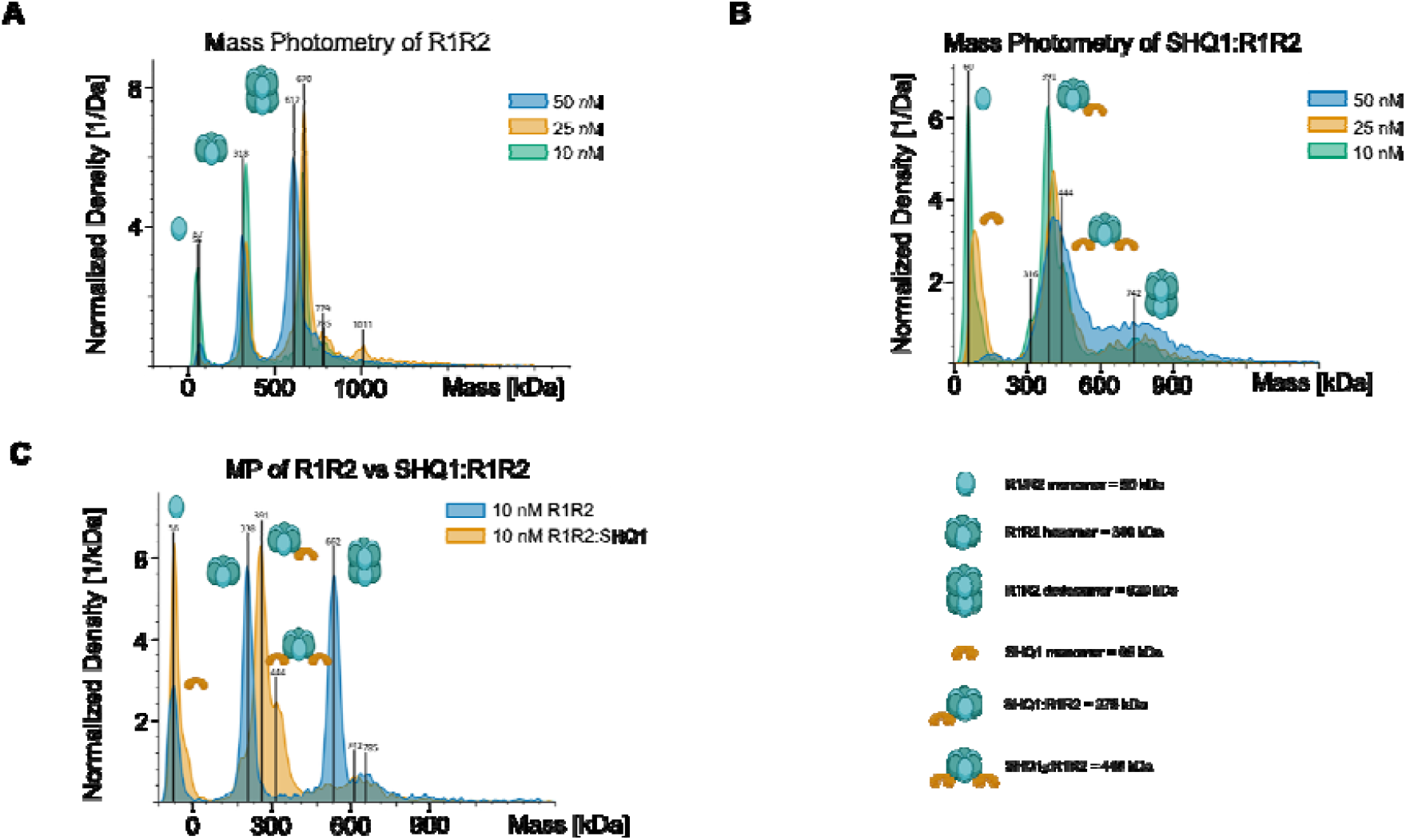
Determination of the R1R2:SHQ1 stochiometry. **A:** Mass Photometry spectrum of isolated R1R2. Three different concentrations (50 [blue], 25 [yellow], and 10 [green] nM) of isolated R1R2 complex were recorded. The spectrum shows three peaks: the smallest molecular weight (59 ± 17 kDa) corresponds to R1 and R2 monomers, which differ slightly in size. The middle peak (338 ± 19kDa) represents R1R2 hexamers, and the highest molecular weight peak (676 ± 21 kDa) embodies the R1R2 dodecamers. At 50 nM (blue) and 25 nM (yellow), the R1R2 complex is mostly in its dodecameric form. Diluting the complexes to 10 nM (green) results in almost the same amount of hexameric and dodecameric forms and increases the formation of R1 and R2 monomers. B: Mass Photometry spectrum of three different concentrations (50 [blue], 25 [yellow], and 10 [green] nM) of the R1R2:SHQ1 complex. The profile shows two distinct peaks: the first at 60 ± 13 kDa and the second at around 384 ± 25 kDa. The first peak represents R1, R2, and SHQ1 monomers; the mass of the second mass peak fits R1R2 hexamers in a complex with a single SHQ1 molecule. The second peak contains a small shoulder at roughly 447 ± 31 kDa, which fits the molecular weight of R1R2 hexamers interacting with two molecules of SHQ1. The 50 nM profile shows the successful complex purification as R1, R2, and SHQ1 monomers, and R1R2 dodecamers are barely detectable. Contrarily, the dilution of R1R2:SHQ1 complexes leads to an increased monomer signal. **C:** Superimposition of Mass Photometry spectra of 10 nM isolated R1R2 complex (blue) and 10 nM R1R2:SHQ1 (yellow). The middle peaks shift toward higher molecular weights when comparing isolated hexameric R1R2 with R1R2:SHQ1. The size shift fits the addition of one molecule of SHQ1 to the R1R2 hexamers. The shoulder of the middle peak in the R1R2:SHQ1 is roughly shifted by the molecular weight of a second SHQ1 molecule. The signal for the R1R2 dodecamer is almost completely abolished in the R1R2:SHQ1 profile.

Nevertheless, dilution may cause the complex to dissociate and allow residual formation of R1R2 dodecamers and monomers (Figure 2B). Figure 2C shows the superimposition of both isolated R1R2 (blue) and R1R2:SHQ1 (yellow) profiles, recorded at 10 nM. The superimposition displays a clear size shift from R1R2 hexamers (blue middle peak) to the largest R1R2:SHQ1 complex species (yellow), which we propose represents the addition of a single SHQ1 molecule to an R1R2 hexamer, thereby reinforcing that SHQ1 disrupts R1R2 dodecamers into hexamers. This peak also displays a shoulder that most likely represents an R1R2 hexamer in complex with two SHQ1 molecules (R1R2:SHQ1_2_), possibly explaining the observed heterogeneity in SPR experiments.

### SHQ1 interacts with the flexible domains II of the R1R2 complex

To map the interaction site between the R1R2 complex and SHQ1, we performed cross-linking mass spectrometry (XL-MS). The purified R1R2:SHQ1 complex was incubated with a 400-fold molar excess of the MS-cleavable cross-linker disuccinimidyl sulfoxide (DSSO) in a test tube. Thus, we used denaturing Mass Photometry as an alternative [42]. Due to the expected mass of the R1R2:SHQ1 complex, traditional SDS-PAGE did not allow for the identification of the oligomeric states stabilized upon DSSO cross-linking. By overlaying the dMP profile of DSSO-cross-linked samples and the native complex profiles, we could show that using 400 molar excesses stabilized all formed complexes. Cross-linking efficiency was assessed by loading the Tris-quenched samples onto an SDS-PAGE. Since all components of the cross-linked complex did not enter the resolving gel but seemed to get retained in the stacking gel (Figure 3A), we concluded that the cross-linking was successful and the concentration of DSSO was sufficient for successful cross-linking.

**Figure 3:**
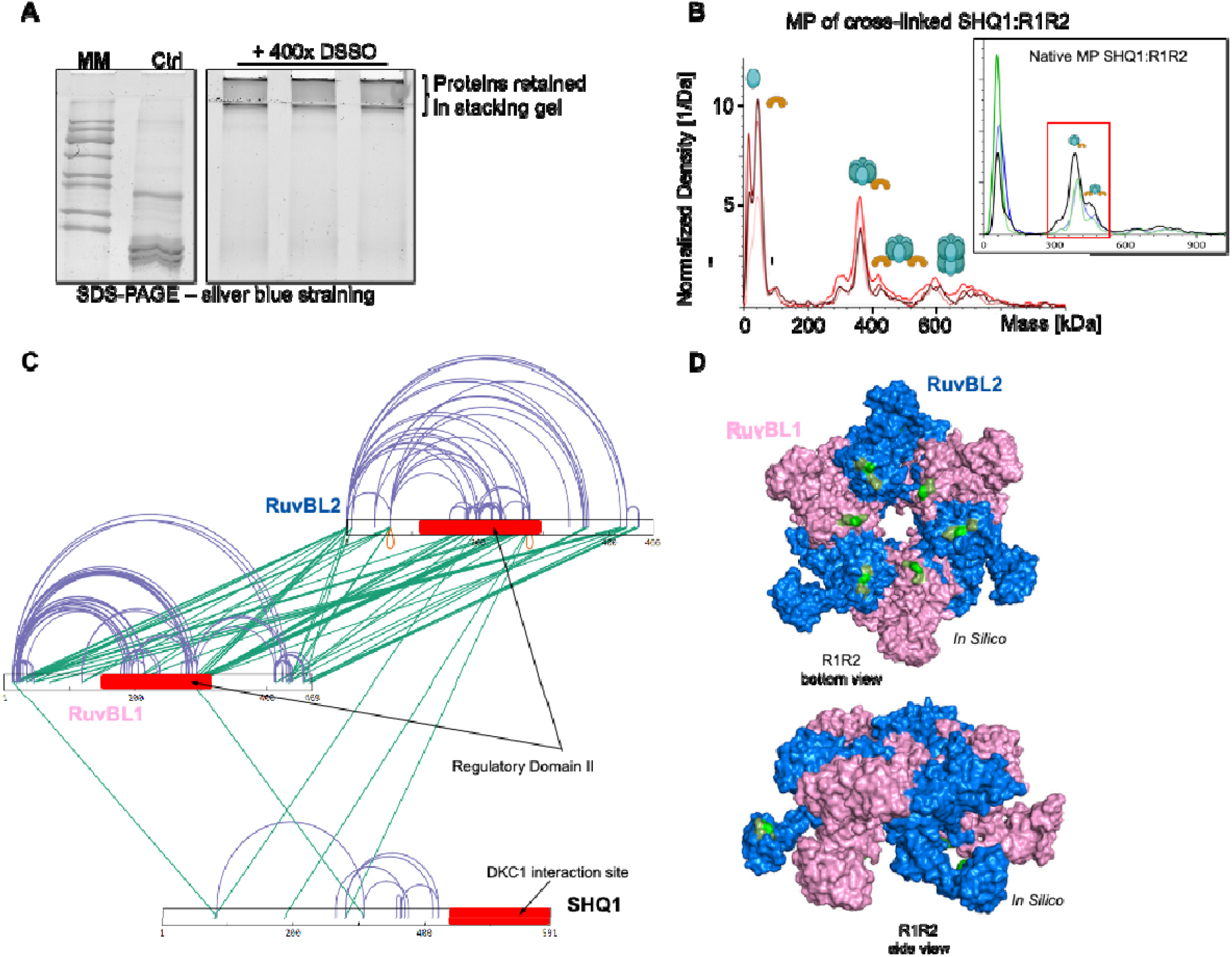
Mapping the interaction site between the RuvBL1:RuvBL2 complex and SHQ1. **A:** The R1R2:SHQ1 complex was loaded to an SDS-PAGE after incubation with 400 times molar excess of the MS cleavable cross-linker DSSO in a test tube. While the control and non-cross-linked sample enters the resolving gel, the cross-linked samples do not enter but accumulate in the stacking gel. **B:** Assessing whether cross-linking introduced artifacts by mass photometry. Mass photometry profiles were recorded in the presence of 8 M urea to guarantee that only the cross-linked complex gave rise to a signal. Comparing the cross-linked profile with the inserted profile of the non-cross-linked R1R2:SHQ1 complex shows that both samples result in comparable profiles. **C:** By mapping the interaction site in cross-linking mass spectrometry experiments, four cross-links between R1R2 and SHQ1 were identified. All cross-links on the R1R2 site were identified in their regulatory DII. For SHQ1, the interactions are clustered on its N-terminal and middle domains. The C-terminal domain is not involved in the interaction with R1R2. **D:** *In silico* model of an R1R2 hexamer with R1 colored in pink and R2 colored in blue. The identified cross-links between R1R2 and SHQ1 (light green) and neighboring polar amino acids (dark green), which may be important for the interaction, are highlighted.

To assess whether cross-linking the complex introduced artifacts, we recorded mass photometry profiles in the presence of 8 M urea, which quenched the signal of non-cross-linked complex [42]. Comparing the cross-linked profiles to those previously recorded for non-cross-linked samples did not reveal any visible differences (Figure 3B). Thus, to identify the complexs’ interaction site, we subjected the DSSO cross-linked R1R2:SHQ1 to cross-linking mass spectrometry (XL-MS). Using XlinkX software, 180 unique cross-links were identified with a 1% False Discovery Rate (FDR). Among the identified cross-links, 60 were intra RuvBL1, 50 were intra RuvBL2, and 7 were intra SHQ1. 62 of the cross-links accounted for inter-molecular RuvBL1/RuvBL2 interactions We mapped 184 inter- and intra-molecular unique cross-links on the published cryo-EM structure of R1R2 (PDB: 7OLE DII conformation open), among which 130 (71 %) were distance validated using a 30 Å Ca-Ca max. threshold. This shows the structural relevance of our dataset and suggests an opened conformation of DII domains. We identified two intermolecular cross-links between SHQ1 and RuvBL1, along with three between SHQ1 and RuvBL2, confirming the presence of R1R2:SHQ1. Interestingly, these cross-links were all within the DII domains of RuvBL1 and RuvBL2, allowing us to position SHQ1 at the level of the DIIs, consistent with previous observations for other RuvBL-client complexes [41, 43].

Table 1 summarizes the interaction sites identified between the R1R2 complex and SHQ1, supporting a DII-mediated mode of association, which is also illustrated in Figure 3C.

**Table 1:** Identified cross-links between the R1R2 complex and SHQ1.

|  |  |
| --- | --- |
| SHQ1 [Y65] | RuvBL2 [K201] |
| SHQ1 [K172] | RuvBL2 [K279] |
| SHQ1 [K265] | RuvBL2 [K279] |
| SHQ1 [K293] | RuvBL1 [K274] |

One of the cross-links identified between SHQ1 and RuvBL1 was located in the linker connecting the His-tag to RuvBL1 (Figure 3C).

Mapping of the R1R2-interacting regions on SHQ1 localizes the interface primarily to its N-terminal and middle domains (Figure 3C). Although the C-terminal domain of SHQ1 was previously proposed to mediate the interaction with R1R2 [35], our XL-MS data on the isolated R1R2:SHQ1 complex does not support its involvement, suggesting that this region remains accessible for interaction with other proteins, such as DKC1 protein factors such as DKC1. Figure 3D highlights the interaction surface of the R1R2 hexamer engaged with SHQ1

For many RuvBL:client protein complexes, flexibility has been reported as an obstacle [43] [44]. To decrease structural dynamism of R1R2:SHQ1 complexes and to determine a high-resolution cryo-EM structure, we performed gradient fixation (GraFix) centrifugation (Figure 4A) [45].

**Figure 4:**
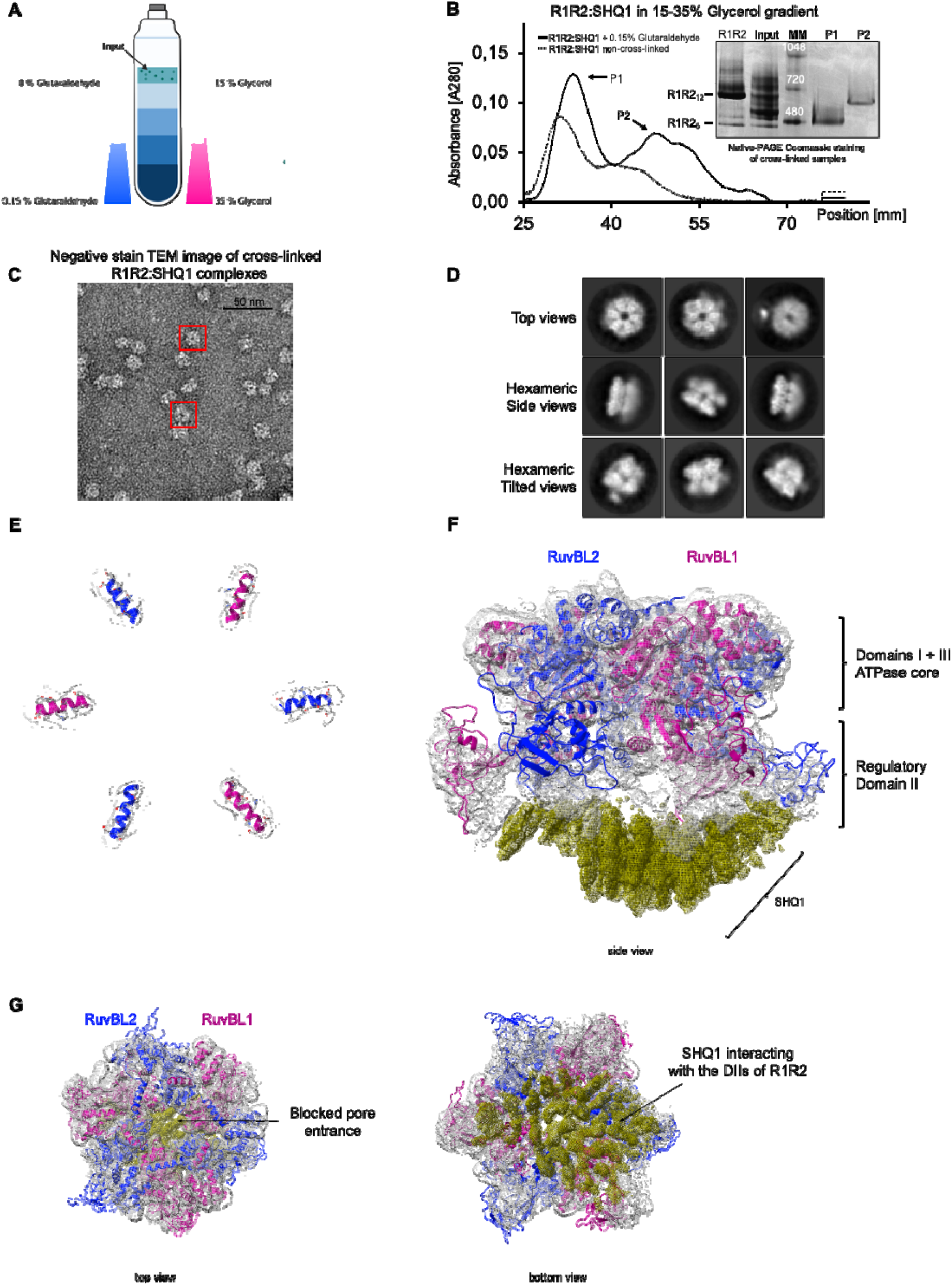
Cross-linking the R1R2:SHQ1 complex improves resolution of 3D reconstruction. **A:** Purified R1R2:SHQ1 complex were loaded to glycerol (15-35 %) and glutaraldehyde (up to 0.15 %) gradients to perform gradient fixation centrifugation (GraFix). While migrating along the gradients, the complexes were mildly cross-linked by increasing concentrations of glutaraldehyde. **B:** Superimposition of GraFixed R1R2:SHQ1 samples in the absence (dashed line) and presence (straight line) of glutaraldehyde. Both profiles display similar characteristics, showing two major peaks: low-density (P1) and higher-density (P2). The inserted Native-PAGE shows the stabilization of P1 at slightly below 480 kDa and P2 closer to 720 kDa. **C:** Negative stain electron microscopy of the low-density peak. The sample shows no signs of aggregation, indicating cross-linking did not introduce artifacts. Additionally, the ring-like structures allow seeing the R1R2 complexes’ characteristic pores. **D:** Selected 2D classes of cross-linked R1R2:SHQ1 complexes. Complexes were detected in top, side, and tilted views. Due to reduced flexibility after cross-linking, the 2D classes seem more detailed than the non-cross-linked classes. No dodecameric R1R2 complexes were detected. **E:** One alpha helix of RuvBL1 (pink) and RuvBL2 (blue) within the AAA+ core module was selected to be represented as ribbons with protruding amino acid side chains represented as sticks, allowing evaluation of the published R1R2 structure (PDB: 7OLE) fitted into our obtained cryo-EM map. **F and G:** Three-dimensional reconstruction of the cross-linked R1R2:SHQ1 complexes. The overall resolution is at 3.9 Å with higher resolution in R1R2s’ DI and DIII and less defined in the DII and SHQ1 area. Compared to the non-cross-linked sample, the reconstruction resulting from cross-linked samples shows increased electron density for SHQ1, which is continuous with the density for the R1R2 hexamer. In top and bottom views, it can be seen that SHQ1 occupies the entrance to the central pore.

Gradients were profiled, which allowed the identification of a lower molecular weight peak (P1) and a higher molecular weight peak (P2) (Figure 4B, continuous line). Both peaks were analyzed by Native-PAGE with subsequent Coomassie staining (Figure 4B insertion), revealing the stabilization of two distinct complex species. Both species were smaller than the R1R2 dodecamers, as comparison with the native R1R2 sample showed (Figure 4B insertion). The higher molecular weight species (P2) migrated between 480 and 720 kDa and is consistent with the size of R1R2:SHQ1_2_. The lower molecular weight species (P1) migrated close to 480 kDa and corresponds to the R1R2:SHQ1 complex. Given its lower apparent molecular weight and expected structural simplicity and homogeneity, the lower molecular weight species was selected as the species of interest, accordingly, fractions corresponding to the center of peak P1 were prepared for 3D structure determination. As a control, R1R2:SHQ1 we also profiled on gradients lacking cross-linking agent, yielding a profile comparable to that obtained under cross-linking conditions (Figure 4B scattered line).

Before loading the complexes onto cryo-EM grids, we first imaged the sample by negative-staining transmission electron microscopy (neg. stain TEM) as a quality control (Figure 4C). This confirmed that the grids were holding particles with characteristic ring-like structures (red box in Figure 4C). Moreover, we did not observe aggregation and observed very little to no dodecameric side views, whose appearance in negative staining EM has been well characterized in previous publications [41]. We then used cryo-EM and single particle analysis to characterize the 3D structure of the cross-linked R1R2:SHQ1 complex. The extracted particles were sorted into 2D classes (Figure 4D), revealing characteristic heterohexameric side and tilted views in which the AAA+ ATPase core module was resolved at high resolution, whereas the protruding flexible DII caused blurring. Top views were also identified, but did not allow distinction between heterohexamers and heterododecamers. The classes representing hexameric views, in addition to all top views, were used to build an *Ab initio* model and were further 3D classified and 3D refined in Cryo-SPARC [46], resulting in a final model of 3.9 Å according to gold standard FSC (Figure 4E). The cryo-EM map allowed the determination of clear secondary-structure elements for the AAA+ core module forming DIs and DIIIs of R1R2s, enabling us to fit a published R1R2 structure (PDB: 7OLE) into the map (Figure 4F). However, we were not able to satisfactorily fit all of the models DIIs into our obtained cryo-EM map, leading us to propose that the conformation of the DIIs involved in SHQ1 binding may differ slightly from the published model. The cryo-EM map shows aa additional continuous electron density below the DIIs (Figure 4E and G). Based on our XL-MS results, we attributed the additional densities placed at the entrance of the R1R2-ring pore to the presence of SHQ1, which was previously reported for other R1R2 clients as well [43, 47]. A precise positioning of SHQ1 relative to the R1R2 hexamer was not possible because of sparse cryo-EM densities of this region of the map, suggesting that although the interaction between SHQ1 alone and the R1R2 hexamer has a very high affinity (0.94 nM), it is highly structurally dynamic.

To confirm that the cross-linking did not impact the integrity of R1R2:SHQ1 complexes, we also performed single particle cryo-EM analysis of non-cross-linked R1R2:SHQ1 complexes (Supplementary Figures 5 and 6). In the resulting cryo-EM map with a resolution of 4.66 Å, we identified an additional density below the R1R2. However, the densities were discontinuous, likely reflecting the high dynamics of the complexes.

### SHQ1 and DKC1 interact simultaneously with the RuvBL1/RuvBL2 complex

Although cross-linking via GraFix enabled detection of a continuous additional density for SHQ1, the resolution did not allow its unambiguous orientation below the RuvBL1/RuvBL2 complex. Hence, we aimed to further rigidify this interaction by exploring the larger physiological complex RuvBL1/RuvBL2/SHQ1 with DKC1. To purify the RuvBL1/RuvBL2/SHQ1/DKC1 complex, of which only DKC1 carried a His-tag at the N-terminus, we co-expressed all four proteins in *E. coli* (pLysS) cells. By capturing DKC1 via its His-tag, we could co-purify all proteins forming a stable complex with it. The eluted sample was subjected to a preparative size-exclusion chromatography column, which showed a large molecular weight peak, possibly representing the RuvBL1/RuvBL2/SHQ1/DKC1 complex isolated from their individual components (Figure 5A, red box). The identity of all four proteins was confirmed by SDS-PAGE (Figure 5B left), by anti-His western blotting identifying DKC1 protein bands (Figure 5B right), and by peptide mapping (Supplementary Figure 7), confirming the co-elution of all four proteins, thus suggesting complex formation. In order to stabilize the complex, they were incubated with a 400 x molar excess of the cross-linking agent DSSO. To monitor the generated stabilized complexes, the sample was loaded on an analytical size-exclusion chromatography column, which showed two distinct peaks of different molecular weights (Figure 5C). While it was possible to isolate the lower-molecular-weight species (Figure 5D, continuous line) by collecting the corresponding fractions, the larger-molecular-weight species remained mixed (Figure 5D, dashed line). By SDS-PAGE, efficient cross-linking was confirmed for both species, as the proteins did not enter the resolving gel of the SDS-PAGE (Figure 5C, insertion). Loading both species on a native PAGE and comparing them with non-cross-linked dodecameric R1R2 complexes (Figure 5D, insertion) revealed that both species show a smeared profile. However, the majority of the lower-molecular-weight species seemed smaller than R1R2 dodecamers, whereas the mixed species appeared mostly larger. The two species were analyzed separately by denaturing mass photometry (Figure 5E-F). The lower-molecular-weight species displayed mostly a single, sharp mass peak at 499 ± 5 kDa, accounting for 85 ± 3% of the total signal, consistent with a homogeneous population. In contrast, the mixed species exhibited two main mass peaks, one at 612 ± 15 kDa (50 ± 6%) and a second at 1085 ± 17 kDa (32 ± 8%). Given its homogeneity and its closer agreement with the calculated theoretical molecular weight of R1R2 in complex with SHQ1 and DKC1 (433 kDa), the lower-molecular-weight species was selected for cryo-EM grid preparation to determine the 3D structure of the R1R2:SHQ1:DKC1 complex.

**Figure 5:**
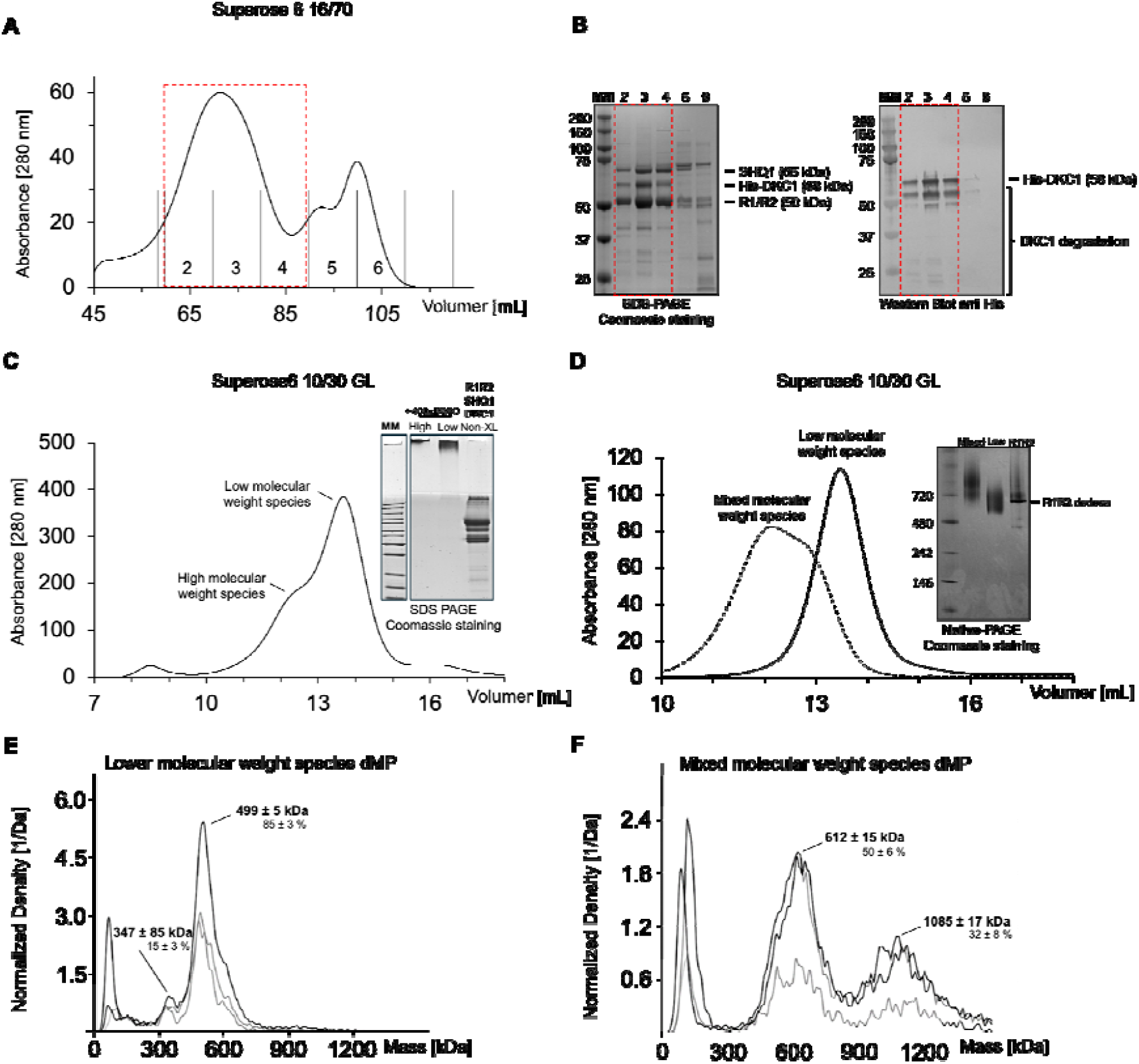
SHQ1 and DKC1 simultaneously interact with the R1R2 complex. **A:** Chromatogram of the eluted proteins after His-tagged DKC1 purification loaded onto a SEC column. The numbers indicate the collected fractions during SEC. The red box indicates the fractions containing the R1R2/SHQ1/DKC1 complex. B: SDS-PAGE (left) and anti-His western blot (right) of the collected fractions resulting from SEC. The red box indicates those fractions highlighted in Figure 5A. Proteins are identified together with their molecular mass on the right side of the gel and western blot. C: Chromatogram of the DSSO cross-linked R1R2/SHQ1/DKC1 complexes loaded to a SEC. The higher- and lower-molecular-weight peaks are identified, and the inserted SDS-PAGE confirms successful cross-linking. D: Chromatograms of the isolated lower molecular weight species (continuous line) and the mixed molecular weight species (dashed line). The inserted Native-PAGE shows that the majority of the lower-molecular-weight species are smaller than R1R2 dodecamer, whereas the mixed-molecular-weight species are mostly larger. E: Denaturing Mass Photometry profile of the lower-molecular-weight species showing 85 ± 3 % of the sample contributes to a mass peak around 499 ± 5 kDa while 15 ± 3 % contributes to a peak around 374 ± 85 kDa. F: Denaturing Mass Photometry profile of the mixed-molecular-weight species showing 50 ± 6 % of the sample contributes to a mass peak around 612 ± 15 kDa, while 32 ± 8 % contributes to a peak around 1085 ± 17 kDa.

### Structural organisation of the RuvBL1/RuvBL2/SHQ1/DKC1 complex at the DII domains

We performed single-particle cryo-EM analysis using the DSSO cross-linked R1R2:SHQ1:DKC1 complex, which enabled the determination of their three-dimensional structure to a refined overall resolution of 6.3 Å (Figure 6A). Albeit the determination of amino acid side chains was not possible, the resolution was sufficient to unambiguously fit RuvBL1, RuvBL2, and, this time, also the AlphaFold predicted structure of SHQ1 into the resulting cryo-EM map using Phenix Predict and build tool. While SHQ1 loop spanning amino acids 293-300 is involved in binding the residues 311 to 314 of DKC1, SHQ1 residues 257-262 form a loop that seems important for the interaction with residues 170-173 and 231-233 of RuvBL2 (Figure 6C). Additionally, the orientation of SHQ1 allowed to propose a binding mode for the AlphaFold predicted structure of DKC1 as well, as the map presented an additional density next to SHQ1. Orienting the SHQ1:DKC1 dimer as previously published for the yeast homologs Shq1:Cbf5 (PDB: 3UAI) (Supplementary Figure 8A) suggests that in humans, DKC1’s PUA domain may interact with SHQ1’s SSD domain, as well (Figure 6B). Our fit also suggests that DKC1 could interact with the counterclockwise-adjacent DII of RuvBL1 (Figure 6D and Supplementary Figure 8D). Although the fit for DKC1 is not confident, the solved structure proves that the presence of DKC1 further rigidifies the interaction site of SHQ1 at R1R2 complexes, thereby improving the resolution corresponding to SHQ1 and allowing its unambiguous fitting.

**Figure 6:**
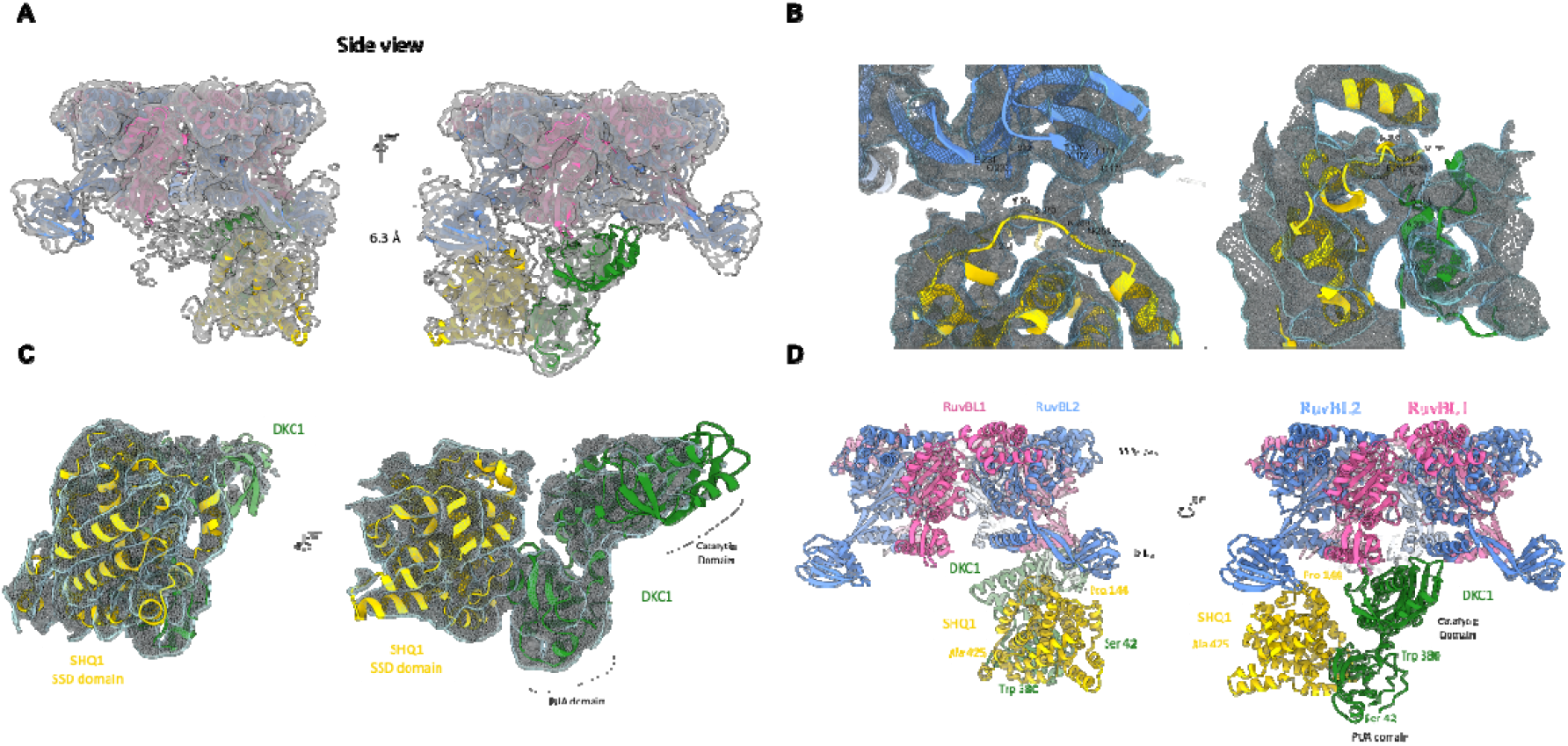
**A:** RuvBL1/RuvBL2/SHQ1/DKC1 model fitted inside the 6.3 Å Cryo-EM map. Phenix Predict and build tool was used to generate starting models of RuvBL1 (pink), RuvBL2 (blue), and a partial SHQ1 (yellow). Partial SHQ1 was then used as a template to position the AlfaFold-generated dimer SHQ1/DKC1. The RuvBL Ring, SHQ1, and DKC1 (green) were further individually fitted using the Flex-EM tool. B: Insight into putative contact regions between RuvBL2 External Domain DII and SHQ1 (left), and SHQ1 and DKC1 (right). Labels are shown for amino acids in close proximity, based on the secondary structure. C: Zoom-in into the density accommodating SHQ1 and DKC1 models. D: Model of RuvBL1/RuvBL2 Hexamer bound to SHQ1:DKC1 dimer. SHQ1 mainly interacts with RuvBL2 (blue) domain II external region, while DKC1 is positioned beneath the neighbour RuvBL1 (pink).

Since both, the DSSO cross-linked R1R2:SHQ1:DKC1 and the GraFix cross-linked R1R2:SHQ1 cryo-EM maps show a density at the entrance to the pore of R1R2 (red density in Supplementary Figure 8B), it strongly suggests that this density corresponds to SHQ1 CS domain. However, as the resolution does not allow the determination of secondary-structure elements, we cannot discard the possibility that this density may correspond to DKC1’s catalytic domain.

It is interesting to note that we identified an additional density in the nucleotide-binding pocket of the RuvBL2 protomer that interacts with SHQ1, which could correspond to either ATP or ADP (Supplementary Figure 8C, left). Since the purification was performed without supplementing nucleotides, and it was impossible to fit three phosphates into the density, we suppose that all ATP molecules might have been hydrolyzed to ADP. For the remaining pockets, no such density could be identified (Supplementary Figure 8C, right). We propose that the binding of SHQ1:DKC1 dimers either rigidifies the nucleotide-binding pocket of RuvBL2, thereby increasing the cryo-EM map’s density to allow the detection of the nucleotide, or this associating heterodimer blocks the dissociation of ADP from RuvBL2. The workflow determining the R1R2:SHQ1:DKC1 structure is depicted in Supplementary Figure 9.

### RPAP3 recruits SHQ1 to RuvBL1:RuvBL2-containing complexes

It has previously been proposed that the R2TP chaperoning complex, along with HSP90, drives the maturation of box H/ACA snoRNPs. To get an understanding of which cellular factors might be involved and to unravel their interplay in the early steps of box H/ACA snoRNP maturation, we fused GFP to SHQ1 and performed a pull-down assay followed by interactome analysis using MS (Figure 7A). Compared with the control (GFP alone), many proteins known to interact with RuvBL1 and RuvBL2 were identified in the SHQ1 pull-down (Table 2). In addition to DKC1, NOP10, NHP2, NAF1, and GAR1, all of which are known to be directly involved in box H/ACA snoRNP maturation, some more unexpected proteins like NOP58, NOPCHAP1, and DPCD were identified.

**Figure 7:**
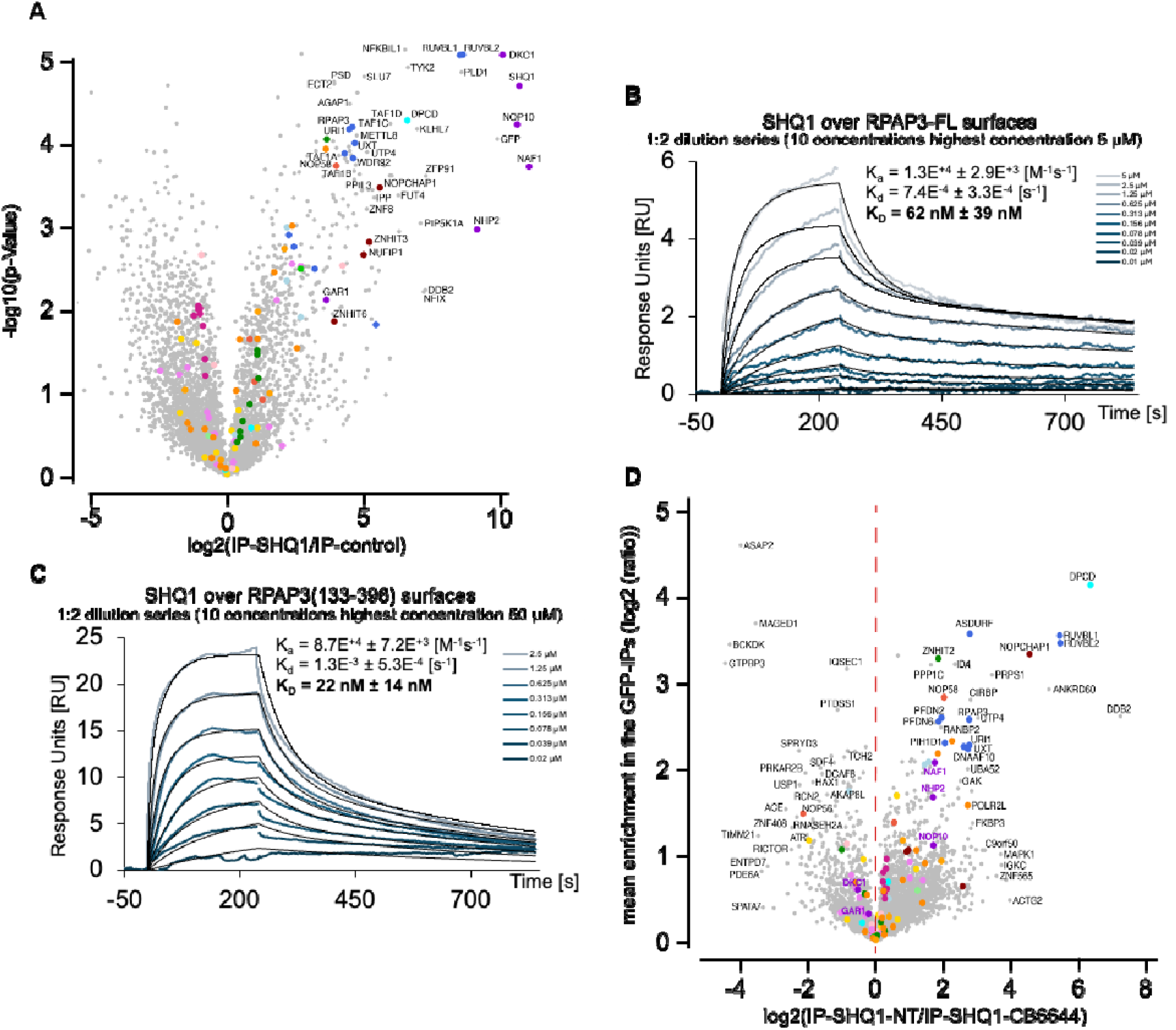
RPAP3 interacts with SHQ1 at the RuvBL1:RuvBL2 complex. **A:** Volcano plot of proteins co-immunoprecipitated with GFP-SHQ1, identified by MS. Parental cells are used as a control. The X-axis represents the log2 of relative protein abundance, while the Y-axis represents the log10 of the students t-test. **B:** Surface Plasmon Resonance sensorgrams of SHQ1 injected into RPAP3-FL surfaces. 10 different concentrations of SHQ1 were generated by 1:2 serial dilutions, with the highest concentration being 5 µM. Fitting the sensorgrams allowed the calculation of the KD for this interaction, which was determined to be 34 nM. **C:** Surface Plasmon Resonance sensorgrams of SHQ1 injected into truncated RPAP3 (133-396) surfaces. 10 different concentrations of SHQ1 were generated by 1:2 serial dilutions, with the highest concentration being 5 µM. Fitting the sensorgrams allowed the calculation of the K_D_ for this interaction, which was determined to be 9.8 nM. **D:** Volcano plot of proteins co-immunoprecipitated with GFP-SHQ1 in presence and absence of CB6644, identified by MS. The X-axis represents the log2 of relative protein abundance of co-purified proteins in non-treated versus treated conditions, while the Y-axis represents the log_2_ of mean enrichment of purified proteins in GFP immunoprecipitations. Proteins that are more immunoprecipitated after CB6644 treatment have a negative sign, while proteins that are more immunoprecipitated in the non-treated cells have a positive sign.

**Table 2:** Proteins identified to be pulled down together with SHQ1. The fold change (FC) for the identified proteins was calculated by dividing the abundance in SHQ1-IP by the abundance in GFP-IP. To simplify the data display in volcano plots, the log_2_ of the FC was calculated. The p-value was calculated to test the statistical significance of the fold changes identified. Values above 2 of the log_10_(p-value) were accepted as statistically significant

| Protein of interest | Fold change SHQ1/Ctrl [$\log_2(\text{FC})$] | Significance [$\log_{10}(\text{p-Value})$] |
| --- | --- | --- |
| **RPAP3** | 4.6 | 4.2 |
| **PIH1D1** | 3.2 | 2.5 |
| **RuvBL1** | 8.5 | 5.1 |
| **RuvBL2** | 8.6 | 5.1 |
| **DKC1** | 10.1 | 5.1 |
| **NOP10** | 10.6 | 4.2 |
| **NHP2** | 9.2 | 3.0 |
| **GAR1** | 3.6 | 2.1 |
| **NAF1** | 11.1 | 3.7 |
| **NOP58** | 4.0 | 3.7 |
| **NOPCHAP1** | 5.6 | 3.5 |
| **DPCD** | 6.6 | 4.3 |

NAF1 has been described as being held in the cytosol upon SHQ1 binding [48]. However, the strong co-purification of DKC1, NOP10, NHP2, RuvBL1, RuvBL2, and NAF1 might indicate an interaction within the nucleus during box H/ACA maturation. For GAR1, we detected only five peptides, which were also less frequently observed than those of other box H/ACA snoRNP components, suggesting that SHQ1 dissociates from the pre-mature complex before the final maturation step, during which NAF1 is exchanged for GAR1 [49, 50].

Regarding the R2TP complex components, RPAP3 was identified with a 24 (log_2_(24.3) = 4.6) times fold enrichment, whereas PIH1D1 was detected considerably less frequently than GAR1, with a fold change of only 9 times. Moreover, PIH1D1 was identified with a confidence only marginally above the acceptance threshold, in contrast to the high confidence with which RPAP3 was detected as an SHQ1 interactor. If both PIH1D1 and RPAP3 were functionally involved in box H/ACA RNP maturation through direct interaction with SHQ1, one would expect both interactions to be robustly represented in the pull-down assays, as observed for RuvBL1 and RuvBL2. The weak and low-confidence detection of PIH1D1, therefore, argues against a prominent or stable interaction with SHQ1 under the tested conditions. In addition, the previously reported interaction interface of PIH1D1 on R1R2 complexes appears to overlap substantially with the SHQ1-binding site on R1R2, suggesting potential competition rather than simultaneous binding [51].

Together, these observations lead us to propose that the early steps of box H/ACA snoRNP maturation may not be driven by the canonical R2TP complex, but rather by an R2T-only assembly lacking PIH1D1. Structural analysis of the RPAP3(281–445):PIH1D1(199–290) complex (PDB: 6GXZ) shows that PIH1D1 engages RPAP3 through its CS domain, contacting residues 281–396 of RPAP3 [47, 52]. Given that SHQ1 also contains a CS domain, we next investigated whether SHQ1 could similarly interact with RPAP3. To this end, purified RPAP3 was immobilized on SPR-compatible CM5 sensor chips, and SHQ1 was injected at ten different concentrations (up to 5 µM) generated by a 1:2 serial dilution. The concentration-dependent signal gain revealed a specific interaction between SHQ1 and RPAP3, with a calculated K_D_ of 34 nM (Figure 7B). To further assess whether SHQ1 and PIH1D1 engage RPAP3 via a similar binding site, we immobilized a truncated form of RPAP3 (residues 133-396), encompassing the previously described PIH1D1 interaction region, and performed SPR analysis using SHQ1 under identical conditions. The calculated K_D_ was approximately 10 nM, displaying binding kinetics comparable to those observed with full-length RPAP3 (Figure 7C). Taken together, these results indicate that SHQ1 and PIH1D1 likely interact with overlapping or identical sites on RPAP3.

Since R1R2 are AAA+ ATPases, ATP hydrolysis may be important during the maturation of box H/ACA snoRNPs. The small molecule CB-6644 is an allosteric inhibitor of the R1R2, locking the complex in a closed/stretched conformation, thereby hindering ATP hydrolysis [53]. We performed SHQ1 pull-downs after treating cells with CB-6644 to get insights into the importance of ATP hydrolysis during early box H/ACA snoRNP biogenesis (Figure 7D) (Table 3).

**Table 3:**
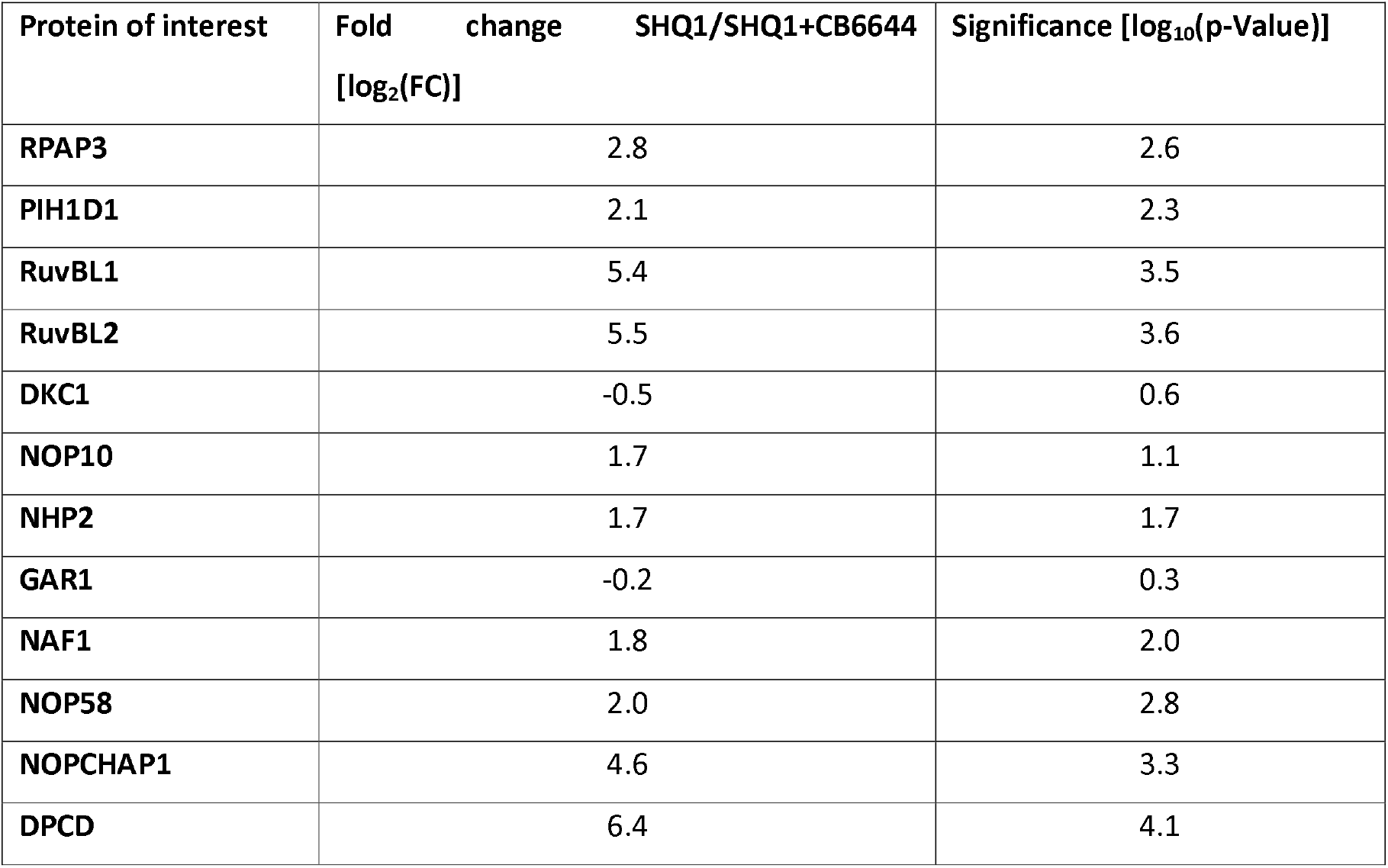
Proteins identified to be pulled down together with SHQ1 in the presence and absence of CB6644. To calculate the FC abundances of proteins found in non-treated control cells were divided by the abundance of the corresponding protein in CB6644 treated cells. Hence, proteins carrying a positive sign are found more frequently in the non-treated control, whereas proteins carrying a negative sign are found more frequently in CB6644-treated cells. The p-value was calculated to test the statistical significance of the fold changes identified.

| Protein of interest | Fold change SHQ1/SHQ1+CB6644<br>[log<sub>2</sub>(FC)] | Significance [log<sub>10</sub>(p-Value)] |
| --- | --- | --- |
| RPAP3 | 2.8 | 2.6 |
| PIH1D1 | 2.1 | 2.3 |
| RuvBL1 | 5.4 | 3.5 |
| RuvBL2 | 5.5 | 3.6 |
| DKC1 | -0.5 | 0.6 |
| NOP10 | 1.7 | 1.1 |
| NHP2 | 1.7 | 1.7 |
| GAR1 | -0.2 | 0.3 |
| NAF1 | 1.8 | 2.0 |
| NOP58 | 2.0 | 2.8 |
| NOPCHAP1 | 4.6 | 3.3 |
| DPCD | 6.4 | 4.1 |

CB6644-treatment resulted in no significant change in SHQ1:DKC1 interaction, suggesting that their interaction is independent of R1R2s’ ATPase activity. However, inhibiting their ATPase activity stabilized the interactions between SHQ1 and RPAP3, as well as with other R2TP-related proteins, among them RuvBL1, RuvBL2, DPCD, URI1, and UXT. Thus, indicating that SHQ1 interacts with RPAP3 in the presence of ATP-loaded R1R2 complexes. The increased association of SHQ1 with R2TP-related proteins in the presence of CB6644 demonstrates that the early steps of box H/ACA snoRNP maturation are most likely linked to the ATPase activity of R1R2, and that their inhibition may also stall the snoRNP maturation.

## Discussion

In this study, we identify SHQ1 as a direct interaction partner of the human RuvBL1:RuvBL2 complex and map the interaction to the regulatory DII domains. Engagement of this region promotes dissociation of the dodecameric R1R2 assembly [41, 54], a mechanism previously described for several R1R2 client proteins [39, 41, 43, 47], suggesting that SHQ1 follows a conserved mode of R1R2 regulation.

Initial cryo-EM analyses were consistent with this model, revealing additional density corresponding to SHQ1 in close proximity to the DII domains of hexameric R1R2 complexes. However, as reported for other R1R2–client assemblies, flexibility associated with the DII domains posed a significant challenge for high-resolution cryo-EM analysis [47]. In line with this, R1R2:SHQ1 complex displayed pronounced internal dynamics, manifested by a dynamic equilibrium between bound and unbound species, heterogeneous 2D class populations, and discontinuous density at the interaction interface. To mitigate these flexibility related limitations and enable structural characterization, we employed gradient fixation centrifugation (GraFix).

Using mass photometry, we confirmed that cross-linking did not introduce artifacts such as higher-order oligomeric species, and 2D classifications of GraFix-treated samples further demonstrated a reduction in the equilibrium between R1R2:SHQ1 complexes and unbound components. This stabilization resulted in a more homogeneous particle population and led to an overall improvement in cryo-EM map quality, particularly in regions corresponding to SHQ1 and the flexible DII domains of RuvBL. In the resulting structure, SHQ1 occupies a position at the entrance of the R1R2-ring pore on the DII side, a region previously implicated in nucleic acid interaction [10, 55]. Given the proposed RNA-mimicking properties of SHQ1 [54, 56], we may speculate that the SHQ1:DKC1 complex remains associated with the hexameric R1R2 chaperone until the arrival of a snoRNA, potentially accessing the complex through the central pore. In this scenario, the snoRNA could either compete with SHQ1 for DKC1 binding [26, 56] or engage DKC1 following an active dissociation of the SHQ1:DKC1 complex, thereby enabling progression of box H/ACA snoRNP maturation.

The improved electron density obtained after GraFix treatment, together with the interaction restraints identified by XL-MS, suggests that SHQ1 engages with more than one DII domain within the R1R2 hexamer ring. Nevertheless, the corresponding density remained insufficient to unambiguously fit the AlphaFold-predicted structure of SHQ1. Incorporation of DKC1 into the R1R2:SHQ1 complex further stabilized the DII region, which clearly appears to constitute the principal interaction platform for both SHQ1 and DKC1. Although the placement of DKC1 within the cryo-EM map remains tentative, the observed density is consistent with a model in which the SSD domain of SHQ1 interacts with the PUA domain of DKC1, as reported for the yeast homologues (PDB: 3UAI). In this configuration, the catalytic domain of DKC1 may additionally be stabilized through contacts with the DII domain of RuvBL1.

At the same time, residues 257-262 within the SSD domain of SHQ1 are positioned to interact with residues 170-173 and 231-233 in the DII domain of RuvBL2, suggesting an interaction mode during which SHQ1 chaperones DKC1’s PUA domain while R1R2 simultaneously chaperones its catalytic domain. The interaction of SHQ1’s SSD domain and the DII of R2 contrasts with our XL-MS data on the R1R2:SHQ1 complex. Hence, we propose that the interaction site between SHQ1 and R1R2 changes in the presence of DKC1, suggesting an active role for R1R2 in segregating the SHQ1:DKC1 dimer.

In line with the increased stabilization of the DII region upon incorporation of the SHQ1:DKC1 heterodimer, we observed additional density in the nucleotide-binding pocket of the RuvBL2 protomer engaged with SHQ1, consistent with the presence of a bound nucleotide. This observation suggests that the interaction of SHQ1 and DKC1 may rigidify the nucleotide-binding pocket of RuvBL2, thereby enhancing local map quality and enabling nucleotide detection. The simultaneous association of SHQ1 and DKC1 with R1R2 hexamers that appear to be at least partially ADP-loaded raises the possibility that ATP binding and/or hydrolysis by the AAA+ ATPases is required for subsequent segregation of the SHQ1:DKC1 complex. Such a mechanism would be consistent with a role for R1R2 in coordinating client release through its ATPase cycle.

Considering that DKC1 has been reported to form dimers within telomerase complexes [57] and that a hetero-tetrameric SHQ1:DKC1 assembly has been isolated from yeast [26], it is plausible that two SHQ1 molecules may simultaneously engage with a single R1R2 hexamer. In this context, the higher molecular weight species separated from the lower molecular weight population in our GraFix and DSSO cross-linking experiments may correspond to biologically relevant R1R2:(SHQ1)_2_ and R1R2:(SHQ1)_2_DKC1_2_ assemblies.

The R2TP complex, an HSP90 co-chaperone composed of hexameric RuvBL1:RuvBL2 associated with RPAP3 and PIH1D1, has previously been proposed to participate in snoRNP assembly [58]. Within this complex, RPAP3 engages the R1R2 hexamer at two distinct sites: first, through its C-terminal domain, which binds the top surface of the rigid AAA+ ATPase core; and second, via an unstructured linker region (residues 420–540) that hangs down the ring and contacts the flexible DII domain [59]. This bipartite interaction mode positions the remaining domains of RPAP3 away from the R1R2 core, thereby leaving them accessible for the recruitment of additional factors. Owing to the symmetry and architecture of the R1R2 ring, up to three RPAP3 molecules can simultaneously associate with a single hexameric R1R2 assembly. In this context, previous in vitro pull-down assays did not detect binding of either RPAP3 or PIH1D1 to SHQ1 [38]. Subsequently, SHQ1 was, however, identified as an RPAP3-associated factor in SILAC-based immunoprecipitation experiments [39], suggesting that this interaction may be transient, indirect, or condition-dependent. Here, we directly demonstrate a specific interaction between SHQ1 and RPAP3 in vitro using surface plasmon resonance, thereby establishing RPAP3 as a bona fide SHQ1-binding partner.

In contrast to RPAP3, PIH1D1 and SHQ1 appear to engage R1R2 hexamers at a similar site, occluding the entrance to the central pore from the DII-facing side of the ring [59, 60]. In light of our observation that R1R2 hexamers can accommodate up to two SHQ1 molecules, potentially together with two DKC1 molecules, the involvement of PIH1D1 in the segregation of SHQ1 and DKC1 from R1R2 appears unlikely, as simultaneous binding at this site would be sterically incompatible. Based on these findings, we propose two alternative, but not mutually exclusive, models for the early steps of box H/ACA snoRNP maturation (Figure 8).

**Figure 8:**
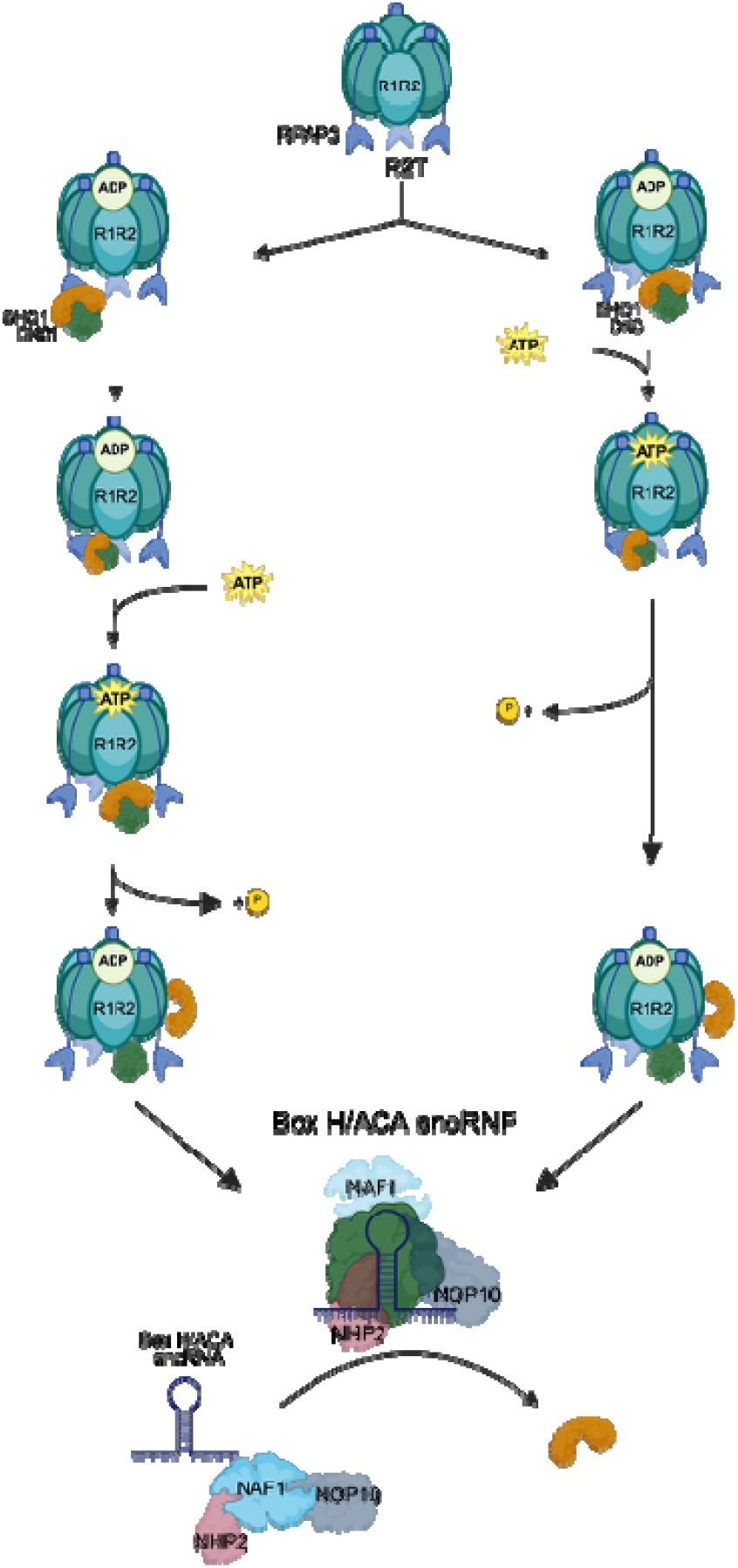
Proposed Mechanism for the early steps of box H/ACA snoRNP maturation. Proposed model of the R2T-assisted early steps of box H/ACA snoRNP maturation. **Left:** RPAP3 recruits the heterodimer of SHQ1:DKC1 by interacting with SHQ1’s CS-domain. Subsequently, RPAP3 hands the SHQ1:DKC1 dimer over to the DII of ADP-loaded R1R2, so that SHQ1 interacts with an R2 protomer, while DKC1 interacts with a R1 protomer. The interaction of SHQ1:DKC1 with R1R2 may induce the nucleotide exchange from ADP to ATP. The ATP hydrolysis-triggered conformational changes of R1R2 might allow the segregation of SHQ1 from DKC1, thereby allowing DKC1 to interact with the box H/ACA snoRNA, NAF1, NOP10, and NHP2 to form another box H/ACA snoRNP intermediate. **Right**: Driven by the nanomolar affinity between SHQ1 and R1R2, the heterodimer of SHQ1:DKC1 interacts with the DIIs of the AAA+ ATPases, which may trigger nucleotide exchange from ADP to ATP. While DKC1 may interact with R1, SHQ1 interacts with R2 and, via its CS-domain, also with RPAP3. ATP hydrolysis and the interaction between RPAP3 and SHQ1 CS-domain may drive the dissociation of SHQ1:DKC1 to allow the formation of the almost mature box H/ACA snoRNP.

In the first model (Figure 8, left), three RPAP3 molecules decorate the RuvBL2 subunits of the R1R2 hexamer, consistent with the known stoichiometry of the R2T complex. Each RPAP3 molecule is capable of recruiting a CS-domain-containing factor, such as SHQ1, PIH1D1, or DPCD [41]. In this scenario, one RPAP3 molecule recruits a SHQ1:DKC1 heterodimer via the CS domain of SHQ1 and subsequently transfers the heterodimer to the DII domains of an ADP-bound R1R2 hexamer. This positioning places SHQ1 in contact with a RuvBL2 protomer, while DKC1 engages a neighbouring RuvBL1 protomer, thereby localizing the heterodimer at the entrance of the central pore of the AAA+ ATPase ring.

In the second model (Figure 8, right), SHQ1:DKC1 heterodimers associate directly with R1R2 hexamers independently of RPAP3. This model is supported by our observation that the R1R2:SHQ1:DKC1 complex is stable in the absence of RPAP3, together with the nanomolar affinity of the individual proteins of SHQ1 and DKC1 [37] for R1R2. In this context, RPAP3 is not required for initial recruitment but may instead play a modulatory role, stabilizing the interaction between SHQ1 and R1R2 and facilitating the subsequent R1R2-mediated dissociation of the SHQ1:DKC1 heterodimer.

In both models, association of the SHQ1:DKC1 complex with R1R2 may promote nucleotide exchange by facilitating ADP release from the AAA+ ATPases, thereby enabling ATP binding and hydrolysis, analogous to a mechanism previously proposed for PIH1D1 [47, 61]. ATP hydrolysis, coupled to conformational rearrangements of the DII domains, and potentially assisted by RPAP3, would then drive the separation of SHQ1 from DKC1. This dissociation would allow DKC1 to engage a box H/ACA snoRNA together with NHP2 and NOP10, promoting formation of a downstream box H/ACA snoRNP intermediate. Following ATP hydrolysis, R1R2 hexamers could be recycled to support subsequent macromolecular assembly reactions involving other CS-domain-containing factors. This framework provides a rationale for how R1R2-containing complexes can participate in the biogenesis of multiple distinct RNP assemblies despite their relatively low cellular abundance [62].

We note that these models may be difficult to reconcile with reports of DKC1 homodimers [22] and SHQ1:DKC1 heterotetramers in yeast [26], as the space beneath the R1R2 DII domains may become sterically crowded. Nevertheless, these higher-order assemblies may represent alternative or transient states that are not simultaneously compatible with R1R2 binding.

Our proposed mechanisms differ from a previously suggested model in which PIH1D1 binds DKC1 and initiates R2TP-mediated recruitment of SHQ1:DKC1 complexes [21]. Future interaction studies dissecting the interplay between R1R2, SHQ1, DKC1, RPAP3, and PIH1D1 will be essential to resolve these alternatives. In particular, while RPAP3 may either mediate recruitment of SHQ1:DKC1 to R1R2 or facilitate their dissociation, PIH1D1 may instead act downstream by recruiting NAF1, which is a target of phosphorylation (contains a FSDDE peptide [63]), and may shuttle NOP10 and NHP2 to the maturation site of box H/ACA snoRNPs [50, 64, 65].

